# Long-term analysis of pertussis vaccine immunity uncovers a memory B cell response to whole cell pertussis immunization that is absent from acellular immunized mice

**DOI:** 10.1101/2021.10.01.462695

**Authors:** Kelly L. Weaver, Catherine B. Blackwood, Alexander M. Horspool, Gage M. Pyles, Emel Sen-Kilic, Emily M. Grayson, Annalisa B. Huckaby, William T. Witt, Megan A. DeJong, M. Allison Wolf, F. Heath Damron, Mariette Barbier

**Affiliations:** West Virginia University, Department of Microbiology, Immunology, and Cell Biology, 64 Medical Center Drive, Morgantown, WV, 26505, USA; Vaccine Development Center at West Virginia University Health Sciences Center, 64 Medical Center Drive, Morgantown, WV, USA

**Author notes:** These authors contributed equally (Kelly L. Weaver, Catherine B. Blackwood).

**Keywords:** *Bordetella pertussis*, whooping cough, vaccine development, whole cell vaccine, vaccination, memory, T_FH_ cells, CXCL13, germinal centers, antigen specific B memory cells, ELISPOTS, outbred mice, pertussis model, acellular vaccine, humoral immunity, immune system, follicular responses, immunological memory, longevity

## Abstract

Over two decades ago acellular pertussis vaccines (aP) replaced whole cell pertussis vaccines (wP) in several countries. Since then, a resurgence in pertussis has been observed, which is hypothesized to be linked to waning immunity. To better understand why waning immunity occurs, we developed a long-term outbred CD1 mouse model to conduct the longest murine pertussis vaccine studies to date, spanning out to 532 days post primary immunization. Vaccine-induced memory results from follicular responses and germinal center formation; therefore, cell populations and cytokines involved with memory were measured alongside protection from challenge. Both aP and wP immunization elicit protection from intranasal challenge and generation of pertussis specific antibody responses in mice. Responses to wP vaccination were characterized by a significant increase in T follicular helper cells in the draining lymph nodes and CXCL13 levels in sera compared to aP mice. In addition, a population of B. pertussis+ memory B cells was found to be unique to wP vaccinated mice. This population peaked post-boost, and was measurable out to day 365 post-vaccination. Anti-B. pertussis and anti-pertussis toxoid antibody secreting cells increased one day after boost and remained high at day 532. The data suggest that follicular responses, and in particular CXCL13 levels in sera, should be monitored in pre-clinical and clinical studies for the development of the next-generation pertussis vaccines.

## INTRODUCTION

Pertussis (whooping cough) is a vaccine-preventable respiratory disease caused by the Gram-negative bacterium *Bordetella pertussis*^1,2^. Whole cell pertussis vaccines (DTP/wP) were first developed and implemented in 1914, but did not become widely available for distribution until the 1940s^1,3^. After their implementation in the United States, DTP vaccines dramatically reduced pertussis disease from ∼200,000 cases a year in the 1930s to ∼2,000 in the 1970s^1,4,5^. However, safety concerns arose that led to the development of acellular pertussis vaccines (aP) in the United States and Europe in the late 1990’s^6–8^. Unlike wPs, which are composed of inactivated *B. pertussis*, aPs contain 1-5 pertussis antigens adsorbed to aluminum hydroxide^1,9^. While aP vaccines protect against symptomatic disease, they do not prevent transmission or asymptomatic carriage of pertussis^10^. Pertussis outbreaks have been increasing at an alarming rate despite high vaccine coverage since wP vaccines were replaced with aP vaccines^11–13^. During the 2012 pertussis outbreak in the US, there were ∼50,000 cases and 20 deaths attributed to this disease, the worst incidence of pertussis observed since the 1950s^14^. Fluctuation of pertussis incidence is hypothesized to be in part due to waning in immunity elicited by the current aP vaccines, with as low as 10% efficacy in between boosters during adolescence^15–17^. The pertussis field has several hypotheses for the resurgence of pertussis including (in no particular order): 1) more sensitive PCR-based testing, 2) increased reporting, 3) bacterial evolution, and 4) lack of duration of the protective immune response^6,18–21^.

Pre-clinical models of pertussis are instrumental to understand vaccine efficacy. Mice have been the primary model used for both experimental vaccine development and to test vaccine lot efficacy. These models allowed evaluation of vaccine efficacy via measurement of immunoglobulin levels and protection against intracranial and intranasal challenge^4,22^. Overall, the murine model has been extremely useful for the study of pertussis, illuminating many aspects of pertussis pathogenesis, immune responses to infection, and vaccine efficacy and development^23–26^. Furthermore, the mouse model has allowed researchers to better understand mucosal responses and neonatal responses to pertussis^27,28^. Recent work has identified several new immune factors associated with pertussis vaccination and intranasal challenge using the mouse model including: T resident memory cells, secretory IgAs, and interleukin-6, to name a few^29–32^. Despite some caveats, such as lack of audible coughing, this model recapitulates disease manifestation, continues to provide novel findings, and remains a relevant model for pertussis vaccine development.

In addition to mice, pertussis vaccine efficacy has been studied in the green olive baboon model, which is currently the gold standard pre-clinical model of pertussis ^18^. The baboon is ideal for studying pertussis vaccines using the human immunization schedule (2, 4, 6, 8 months) with subsequent challenge at 9 to 12 months^10,33^. At this age, baboons cough and show similar disease manifestation as humans. The baboon model showed that aP immunized hosts are able to be infected and transmit *B. pertussis*, which contributes to the current pertussis problems^34–36^. Unfortunately, baboons are not suitable for longevity studies because after 15 months of age, they become resistant to infection (Tod Merkel; personal communication). Other models of pertussis have been developed, including the rat model that allows for measurement of the hallmark cough of pertussis, and mini pigs that were recently used to study longevity of pertussis vaccines^37,38^. While highly useful to study pertussis and vaccine-mediated responses, these models are still under-developed and suffer from the lack of broadly available tools and reagents. In the murine model, immunological memory responses are well characterized and there are a variety of established tools available to study vaccine-induced memory. Therefore, in this study, we focused on adapting the mouse model of pertussis to study the longevity of the pertussis vaccine-mediated humoral memory response.

Immunological memory is the ability of the immune system to rapidly recognize and respond to an antigen after prior exposure^39^. In order to induce immunological memory, B cell affinity maturation reactions, known as follicular immune responses, occur in microenvironments known as germinal centers (GCs)^16^. GCs are located within the draining lymph node near the site of vaccination, and in the spleen^18^. T follicular helper cells (T_FH_ cells) are crucial for the formation of GCs, B cell affinity maturation, development of memory B cells (MBCs), and high affinity antibody secreting cells^40–43^. T_FH_ cells and follicular dendritic cells in the GC express a signaling molecule known as C-X-C motif chemokine ligand 13 (CXCL13) which binds to C-X-C motif chemokine receptor 5 (CXCR5). CXCR5 is a G-protein coupled receptor expressed in T_FH_ cells and B cells^44^. CXCL13:CXCR5 interactions play a role in recruitment of B cells to the follicle and in the organization of the GC^45,46^. Once B cells have migrated to the follicle in response to CXCL13, high affinity B cells are selected by T_FH_ cells to migrate from the light zone to the dark zone where they undergo proliferation and somatic hypermutation. The products of the GC reaction are MBCs and antibody secreting plasma cells that function to persist overtime and orchestrate the elimination of a pathogen if it is encountered.

The objective of this study was to examine the follicular responses induced by vaccination against pertussis in mice to gain insights into the duration of vaccine-induced memory. To do so, we developed a long-term murine model of pertussis vaccine prime, boost, and intranasal challenge. We used this model to compare aP and wP protection and follicular responses beginning at day 20 and concluding at day 532 post-prime. Given that administration of wP leads to longer-lasting protection than aP in humans, we hypothesized that wP would induce more robust follicular responses than aP immunization in mice^47^. Responses to vaccination were measured by quantification of pertussis-specific antibody titers, antibody secreting cells, and follicular responses. Protection provided by vaccination was measured by quantifying bacterial burden in the airways after challenge. In this work, we describe immunological memory markers that are significantly increased in wP and not aP immunization, such as CXCL13 and antigen-specific memory B cell production. Our data identify that B memory responses are an underappreciated aspect of pertussis immunity that could guide future pertussis vaccine development.

## MATERIALS AND METHODS

### B. pertussis strains and growth conditions

*Bordetella pertussis* strain UT25Sm1 was kindly provided by Dr. Sandra Armstrong (University of Minnesota). UT25Sm1 strain has been fully genome sequenced (NCBI Reference Sequence: NZ_CP015771.1). UT25Sm1 was grown on Remel Bordet Gengou (BG) agar (Thermo Scientific, Cat. #R452432) supplemented with 15% defibrillated sheep blood (Hemostat Laboratories, Cat. #DSB500) and streptomycin 100 μg/mL (Gibco™, Cat. #11860038) at 36°C for 48 hours. Bacteria were then collected using polyester swabs and resuspended in Stainer Scholte media^48^ (SSM) supplemented with L-proline and SSM supplement. SSM liquid culture was incubated for 24 hours at 36°C with constant shaking at 180 rpm until reaching OD_600nm_ 0.5 with 1 cm path width (Beckman Coulter™ DU 530 UV Vis spectrophotometer). The UT25Sm1 *B. pertussis* culture was diluted in supplemented SSM to OD_600nm_ = 0.24 - 0.245 (equivalent to 10^9^ CFU/mL) to be used for challenge or serological analysis by ELISA.

### Vaccine preparation and immunization, bacterial challenge, and euthanasia

The World Health Organization (WHO) standard whole cell *B. pertussis* vaccine (wP) was obtained from the National Institute for Biomedical Standards and Control (NIBSC, Cat. #94/532, batch 41S) and compared to the acellular *B. pertussis* vaccine DTaP (Infranrix®, GlaxoSmithKline). It is important to note that the NIBSC wP is not a DTP alum adjuvanted human vaccine. Both wP and DTaP vaccines were diluted to 1/10^th^ the human dose. Phosphate Buffered Saline (PBS) (Millipore Sigma™, Cat. #TMS012A) was administered as a vehicle control. Vaccine injections were administered at a volume of 50 μL intramuscularly in the right hind limb. In all experimental groups, 6-week-old outbred female CD1 mice (Charles River, Strain code 022) were used. Mice were intramuscularly primed at day 0, followed by a booster of the same vaccine at day 21. Mice were euthanized at days 20, 22, 35, 60, 90, 152, 365, and 532 post-vaccination. Three days prior to euthanasia, mice were anesthetized by intraperitoneal injection (IP) ketamine (7.7 mg/kg) (Patterson Veterinary, Cat. #07-803-6637) and xylazine (0.77 mg/kg) (Patterson Veterinary, Cat. #07-808-1939) in sterile 0.9% NaCl (Baxter, Cat. #2F7124) and challenged intranasally with ∼2×10^7^ CFU/dose of live *B. pertussis*. At day three post-challenge mice were euthanized by intraperitoneal injection of Euthasol (390 mg pentobarbital/kg) (Patterson Veterinary, Cat. #07-805-9296) in sterile 0.9% *w/v* NaCl.

### Quantification of bacterial burden

Lung and trachea homogenates as well as nasal lavage (nasal wash) were collected post mortem and used to enumerate bacterial burden per tissue. Mice were challenged at days 38, 63, 93, 155, 368, and 535 post-prime. The nasal cavity was flushed with 1 mL sterile PBS for nasal lavage. The lung and trachea were homogenized separately in 1 mL sterile PBS using a Polytron PT 2500 E homogenizer (Kinematica). Samples were serially diluted in ten-fold dilutions in PBS and plated on BG agar to quantify viable bacterial burden. Plates were incubated at 36°C for 48 hours to determine colony forming units (CFUs) per mL.

### Serological analysis of immunized mice

Enzyme linked immunosorbent assay (ELISA) was utilized to measure antigen-specific antibodies in the serum of immunized mice^2,49–51^. After euthanasia, blood was collected in BD Microtainer serum separator tubes (BD, Cat. #365967) via cardiac puncture at days 20, 22, 35, 60, 90, 152, 365, and 532 post primary immunization. Blood was centrifuged at 14,000 *x g* for 2 minutes and sera were stored at -80°C. Pierce™ high-binding 96 well plates (Thermo Scientific™, Cat. #15041) were coated with 5×10^7^ CFU/well viable *B. pertussis*, 6.25 ng/well of diphtheria toxoid (List Biological Laboratories, Cat. #151), 6.25 ng/well tetanus toxoid (List Biological Laboratories, Cat. #191), 50 ng/well pertussis toxin (List Biological Laboratories, Cat. #180) or 50 ng/well of filamentous hemagglutinin (Life Sciences, Inc., Cat. #ALX-630-123-0100 Enzo) overnight at 4°C. Plates were washed three times with PBS-Tween 20 (Fisher Scientific, Cat. #BP337-500), then blocked with 5% *w/v* non-fat dry milk (Nestle Carnation, Cat. #000500002292840) in PBS-Tween 20. Serum samples were diluted 1:50 using 5% *w/v* milk in PBS-Tween 20 and serially diluted to 1:819,200 for anti-*B. pertussis* and anti-FHA ELISAs. Serum samples were diluted 1:200 using 5% *w/v* milk in PBS-Tween 20 and serially diluted to 1:3,276,800 for anti-pertussis toxin, anti-diphtheria toxoid, and anti-tetanus toxoid ELISAs. Plates were incubated at 37°C for 2 hours and washed four times with PBS-Tween 20. Secondary goat anti-mouse IgG antibody 1:2000 (Southern Biotech, Cat. #1030-04) conjugated to alkaline phosphatase was added and incubated for 1 hour at 37°C. Wells were washed five times with PBS-Tween 20 and Pierce p-Nitrophenyl Phosphate (PNPP) (Thermo Scientific, Cat. #37620) was added to each well to develop plates for 30 minutes in the dark at room temperature. The absorbance at 405 nm was read utilizing a SpectraMax^®^ i3 plate reader (Molecular Devices). The lower limit of detection for serum titers was 1:50, and for statistical analysis, all values below the limit of detection are represented with the arbitrary value of one. Endpoint titers were determined by selecting the dilution at which the absorbance was greater than or equal to twice that of the negative control.

### Tissue isolation, preparation, staining, and flow cytometry

Flow cytometry was used to characterize cell populations from the spleen and inguinal lymph nodes. Organs were harvested at days 20, 22, 35, 60, 90, 152, 365, and 532 post-prime. Spleen and lymph nodes were homogenized using disposable pestles (USA Scientific, Cat. #1405-4390) in Dulbecco’s Modified Eagle Media (DMEM) (Corning Incorporated, Cat. #10-013-CV) with 10% *v/v* fetal bovine serum (FBS) (Gemini Bio, Cat. #100-500). Homogenized samples were strained for separation using 70 μM pore nylon mesh (Elko Filtering Co, Cat. #03-70/33) and centrifuged for 5 minutes at 1,000 *x g*. Splenocytes were resuspended in 1 mL red blood cell lysis buffer BD Pharm Lyse (BD Biosciences, Cat. #555899) and incubated at 37°C for 2 minutes. After red blood cell lysis, samples were centrifuged 1000 *x g* for 5 minutes and resuspended in PBS with 5mM Ethylenediaminetetraacetic acid (EDTA) (Fisher Scientific, Cat. #50-103-5745) and 1% *v/v* FBS. Single cell suspensions were incubated with 5 μg/mL anti-mouse CD16/CD32 Fc block (clone 2.4G2, Thermo Fisher Scientific, Cat. #553142) for 15 minutes at 4°C per the manufacturer’s instructions. Cells from tissues were stained with antibodies against cell surface markers (**S2 Table**). Each single cell suspension was incubated with the antibody cocktails for 1 hour at 4°C in the dark. Samples were washed by resuspending in PBS, centrifuging, removing the supernatant, and washing in PBS with 5mM EDTA and 1% *v/v* FBS and fixed with 0.4% *w/v* paraformaldehyde (Santa Cruz Biotechnology, Cat. #sc-281692) overnight. After fixation, samples were centrifuged and washed before resuspension in PBS with 5mM EDTA and 1% *v/v* FBS. The samples were processed using an LSR Fortessa flow cytometer (BD Biosciences) and analyzed using FlowJo (FlowJo™ Software Version v10). Cells were counted using Sphero AccuCount 5-5.9 μm beads according to the manufacturers protocol (Spherotech, Cat. #ACBP-50-10).

### Bacterial preparation for antigen-specific memory B cell purification

*B. pertussis* was grown as described above. SSM liquid culture was diluted to 10^6^ CFU/mL using PBS, aliquoted into 1.5 mL tubes, and inactivated by heating at 65°C for 1.5 hours with constant shaking. Heat-killed bacteria were then stained with BacLight Red (Invitrogen™, Cat. #B35001) overnight per the manufacturer’s instructions. The fluorescently labeled *B. pertussis* cells were centrifuged at 15,000 *x g* for 15 minutes, supernatant was removed, and the labeled *B. pertussis* cells were resuspended in DMEM with 10% *v/v* FBS for incubation with splenocytes from immunized mice.

### Antigen-specific memory B cell purification

Spleens were extracted into DMEM + 10% *v/v* FBS and homogenized with a small pestle and centrifuged at 1000 *x g* for 5 minutes. Cells were resuspended in DMEM with 10% *v/v* FBS and filtered through 70 μM pore nylon mesh. Filtrate was centrifuged at 1000 *x g* for 5 minutes. Cell pellets were resuspended in 1 mL BD Pharm Lyse (BD Biosciences, Cat. #555899) at 37°C for 3 minutes to lyse red blood cells. The remaining cells were centrifuged at 1000 *x g* for 5 minutes at 4°C. The supernatant was discarded, and the pellet was resuspended in DMEM with 10% *v/v* FBS. The remaining cells were centrifuged at 1000 *x g* for 5 minutes at room temperature and resuspended in 6×10^7^ CFU/mL heat-killed fluorescently labeled *B. pertussis* reconstituted in DMEM with *v/v* 10% FBS. Splenocytes and fluorescently labeled *B. pertussis* were mixed end over end for 1 hour at 4°C.

After incubation, the cells were centrifuged at 1000 *x g* for 5 minutes and resuspended in a cocktail containing the Miltenyi Memory B Cell Biotin-Antibody Cocktail (Miltenyi Memory B Cell Isolation Kit, Cat. #130-095-838), Miltenyi anti-IgG1-APC and Miltenyi anti-IgG2-APC antibodies for a total volume of 500 µL per sample. Cells and fluorescently labeled bacteria were mixed using an end over end rotator for 5 minutes at 4°C. After incubation, anti-biotin magnetic beads (Miltenyi Memory B Cell Isolation Kit) were added to the cocktail. The samples were mixed end over end in the dark for 10 minutes at 4°C. Samples were then transferred to 15 mL conical tubes and non-memory B cells were depleted using the AutoMACS Magnetic Sorter (Miltenyi Biotec) “deplete” program. The negative fraction was retained and centrifuged at 1000 *x g* for 5 minutes. Samples were resuspended with anti-APC magnetic beads (Miltenyi Memory B Cell Isolation Kit), and mixed end over end in the dark for 15 minutes at 4°C. PBS with 1% *v/v* FBS and 5 mM EDTA was added to the samples before centrifuging at 1000 *x g* for 5 minutes. Supernatant was discarded and samples were resuspended in PBS + 1% FBS + 5 mM EDTA. Memory B cells were enriched using the AutoMACS “possel_s” program. The positive fraction was retained, and the cells centrifuged at 1000 *x g* for 5 minutes. Supernatant was discarded and the memory B cells resuspended in 1 mL PBS with 1% *v/v* FBS and 5 mM EDTA. Memory B cells were diluted to 10^6^ cells/mL and resuspended in 100 µL of antibody cocktail (**S2 Table B)** for further memory B cell analysis by flow cytometry.

### ELISpot sample preparation and analysis

The Mouse IgG ELISpot (ImmunoSpot®, Cat. #mIgGIgA-DCE-1M/10) was utilized to quantify antibody secreting cells specific for *B. pertussis*. UT25Sm1 was cultured as described above. PVDF membrane 96-well plates were coated with 5×10^7^ CFU/well *B. pertussis* and incubated overnight at 4°C. To measure pertussis toxoid specific antibody secreting cells, wells were coated with 50 ng/well heat inactivated pertussis toxin. Bone marrow samples were isolated by centrifuging hind femurs at 400 *x g* for 5 minutes in 200 µL PCR tubes with holes in the bottom that were placed into 2 mL Eppendorf tubes. The bone marrow was resuspended in heat-inactivated filter-sterilized FBS and filtered through 70 µm mesh with FBS with 10% *v/v* dimethyl sulfoxide (DMSO) (Sigma-Aldrich, Cat. #D8418-100ML) and stored at -80°C. To run the assay, cells were thawed in a 37°C water bath and placed in DMEM with 10% *v/v* FBS. Cells were centrifuged at 400 *x g* for 5 minutes, resuspended in CTL Test B Culture medium (ImmunoSpot), diluted 1:10 with PBS and 1:1 with trypan blue stain (Invitrogen™, Cat. #T10282), and counted on the Countess II Automated Cell Counter (Invitrogen). Plates were washed with PBS and bone marrow cells were added to the first row then serially diluted two-fold down the plate. Cells were incubated at 36°C overnight and then imaged and counted using the ImmunoSpot® S6 ENTRY Analyzer and CTL Software. Dilutions with spots ranging from ∼10-100 per well were selected to enumerate the number of anti-*B. pertussis* antibody-producing cells per sample. Cell counts were normalized to spots per 10^6^ cells using the cell and spot counts.

### Statistics

Statistical analysis was performed using GraphPad Prism version 8 (GraphPad). To compare two groups an unpaired Student’s *t*-test was used. When comparing three or more groups of parametric data a one-way ANOVA (analysis of variance) with Tukey’s multiple comparison test was used unless otherwise noted. For non-parametric data a Kruskal-Wallis test with Dunnet’s post-hoc test was used.

### Animal care and use

All mouse experiments were approved by the West Virginia University Institutional Animal Care and Use Committees (WVU-AUCU protocol 1901021039) and completed in strict accordance of the National Institutes of Health Guide for the care and use of laboratory animals. All work was done using universal precautions at BSL2 under the IBC protocol # 17-11-01.

## RESULTS

### Use of a long-term vaccine memory model using outbred CD1 mice

Immune memory against pertussis varies greatly depending on the vaccine used. It is estimated that the duration of protection conferred by wP vaccines lasts 4-12 years^52^. For aP vaccines, immunity wanes much more quickly, as observed and underscored by the outbreaks in California and other locations^53^. In our study, we set out to establish a long-term pertussis vaccine efficacy model to evaluate the duration of immunity and identify additional factors that contribute to either wP efficacy or the inadequate responses of aPs. To model the human outbred population, we selected outbred CD1 mice that are also known to have a long lifespan. We aimed to compare the DTaP vaccine (Infanrix; GSK) to a prototype whole cell pertussis vaccine. DTP vaccines are no longer available in the US; therefore, we used the NIBSC whole cell pertussis antigen vaccine for comparison. Unlike most combination DTP vaccines, the NIBSC whole cell pertussis vaccine is not adjuvanted with alum nor combined with diphtheria and tetanus toxins. Female CD1 mice were mock-immunized (PBS), immunized with 1/10^th^ human dose DTaP, or 1/10^th^ human dose of NIBSC wP vaccine at six weeks of age by intramuscular administration and subsequently boosted three weeks later. We performed our analysis on mice at day 20 post prime (1 day pre-boost), day 22 (1 day post-boost), day 35 (2 weeks post-boost; *Bp* challenge), day 60 (∼5.5 weeks post-boost; *Bp* challenge), day 90 (∼10 weeks post-boost; *Bp* challenge), day 152 (∼19 weeks post-boost; *Bp* challenge), day 365 (∼49 weeks post-boost; *Bp* challenge), and at day 532 (∼73 weeks post-boost; *Bp* challenge) (**Fig 1A**).

**Fig 1.**
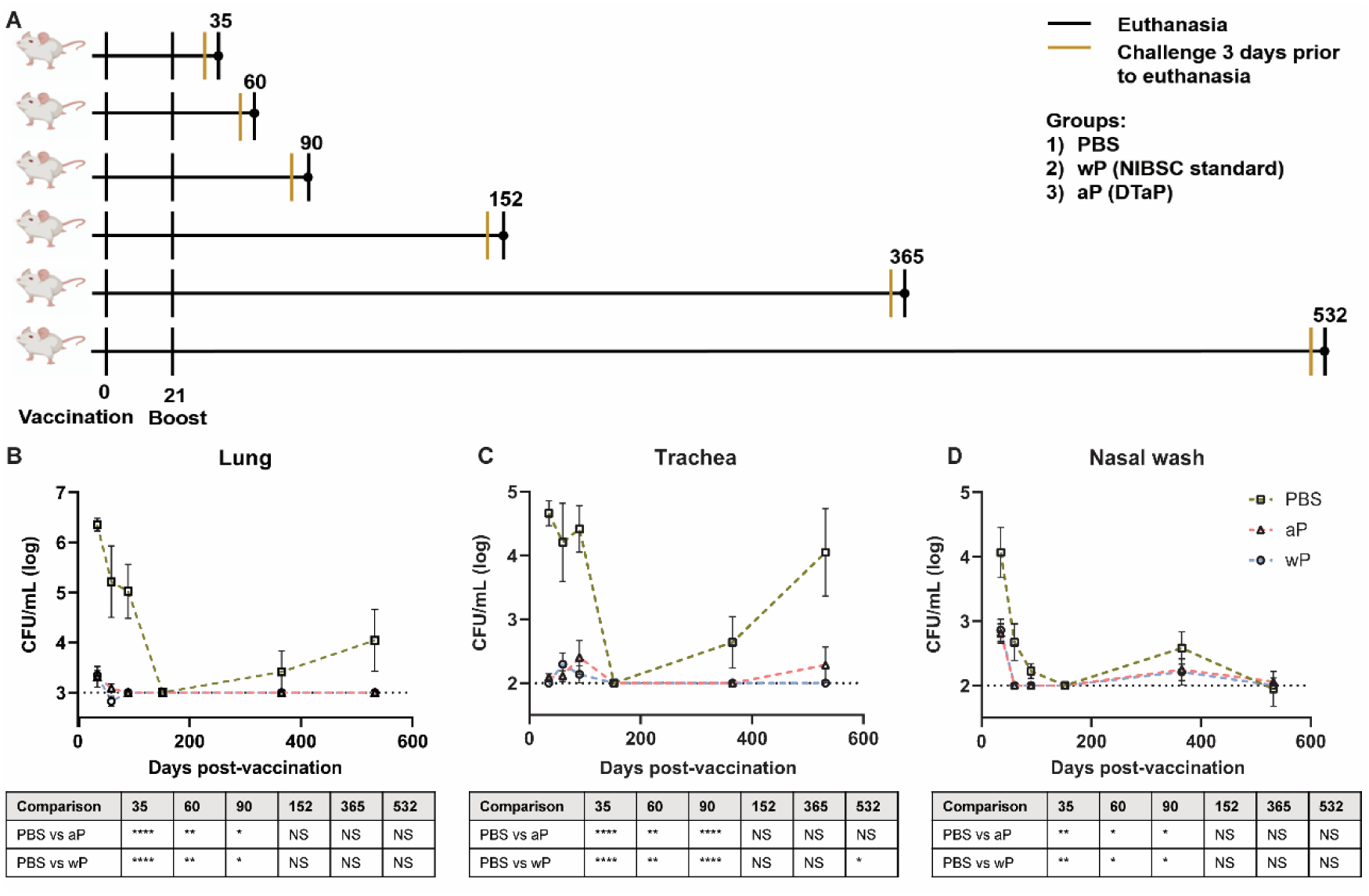
Susceptibility to *B. pertussis* in CD1 mice changes depending on age. **A)** Experimental design and vaccination timeline. Mice were vaccinated at day 0 and boosted at day twenty-one. Vertical black lines indicate the day that non-challenged mice were processed post-vaccination. Mustard colored lines represent groups of mice that were challenged three days prior to processing. **B, C, D)** Bacterial burden in mice challenged with *B. pertussis*. Mice were vaccinated on day 0 with PBS, aP, or wP at 1/10^th^ the human dose and boosted with the same vaccine at day 21. Mice were challenged with 2×10^7^ CFU/dose by intranasal instillation 3 days prior to euthanasia. Bacterial burden of PBS (n=4-16), aP (n=4-16), and wP (n=4-16) vaccinated mice in the lung **(B)**, trachea **(C)**, and nasal wash **(D)**. PBS was used as a vehicle control. Data represent one to four independent experiments. Data were transformed using Y=Log(Y). The *p*-values were calculated using ANOVA followed by a Tukey’s multiple-comparison test, \**p* < 0.05, \*\**p* < 0.01, and \*\*\*\**p* < 0.0001. Error bars are mean ± SEM values.

### wP and aP vaccines provide long-term protection in CD1 mice against respiratory challenge with B. pertussis

To design the next generation of pertussis vaccines, it is important to understand the underlying immunological cause of the relatively short-term immunity provided by aP vaccines. aP vaccines were originally developed and tested in mouse models that provided limited information regarding the longevity of protection. Furthermore, it remains to be determined if this model can mimic the waning immunity observed in humans. We hypothesized that mice immunized with 1/10^th^ the human dose of wP and aP would be protected from challenge early on, but that protection would decrease over time in aP immunized mice. Mice were intranasally challenged with 2×10^7^ CFU/dose of *B. pertussis* at multiple time points between day 35 and day 532 post-vaccination. Mice were then euthanized three days post-infection to study vaccine-mediated protection. The bacterial burdens in the lung (**Fig 1B**), trachea (**Fig 1C**), and nasal wash (**Fig 1D**) were determined by plating serial dilutions and colony counting. Mice vaccinated with vehicle control (PBS) and intranasally challenged at days 35, 60, and 90 post-vaccination had high bacterial burden in the lung, trachea, and nasal wash. Surprisingly, vehicle control immunized mice challenged at day 152 post-vaccination had undetectable bacterial burden in the airways. By days 365 and 532 post-vaccination, bacterial burden was again detectable in the vehicle control mice. When comparing the efficacy of aP and wP vaccines over time, we observed that bacterial burden in the lung (**Fig 1B**), trachea (**Fig 1C**), and nasal wash (**Fig 1D**) was significantly decreased in immunized mice, regardless of the vaccine administered, compared to the vehicle control group at days 35, 60, and 90. Protection remained significant at day 532 in the trachea of wP immunized animals compared to mock-vaccinated animals (**Fig 1D**). At this dose, both aP and wP vaccines protected mice from intranasal challenge. The data illustrate unique susceptibility of CD1 mice to *B. pertussis* over time and the importance of longitudinal studies to identify the optimal timeframe to study vaccine efficacy.

### Pertussis specific antibody responses to aP and wP immunization persist as far as day 532 post-prime

To gain insights into the differences between aP and wP immunological responses, the model described above was applied to study pre- and post-boost responses, and their evolution over time. The subsequent studies were performed in non-challenged animals to clearly separate response to vaccination from response to challenge.

Antibodies play a major role in vaccine-mediated protection against numerous pathogens and are a correlate of protection used to evaluate or predict vaccine efficacy^6,54^. DTP/wP and DTaP vaccines induce opsonizing antibodies that contribute to bacterial clearance and induce anti-pertussis toxin (PT) antibodies that neutralize toxin function^55,56^. However, the value of some of these antibodies in protection against pertussis disease or *B. pertussis* infection is highly debated^57^. Here, we hypothesized that antibody responses to *B. pertussis* antigens in aP immunized mice would decrease over time compared to wP immunized mice. We first measured anti-*B. pertussis* and anti-FHA IgG antibodies in the serum of immunized mice over time to determine the levels of surface-binding antibodies (**Fig 2A, 2B**). Both aP and wP immunization elicited significant production of anti-*B. pertussis* and anti-FHA IgG antibodies compared to vehicle-control immunized mice (**Table S1**). Anti-*B. pertussis* antibody levels were not statistically different between aP and wP vaccinated groups, and they peaked after boost and remained elevated out to day 532 post-prime (**Fig 2A, Table S1**). Anti-FHA antibodies also increased after boost and remained elevated at day 532 post-prime (**Fig 2B**).

**Figure 2.**
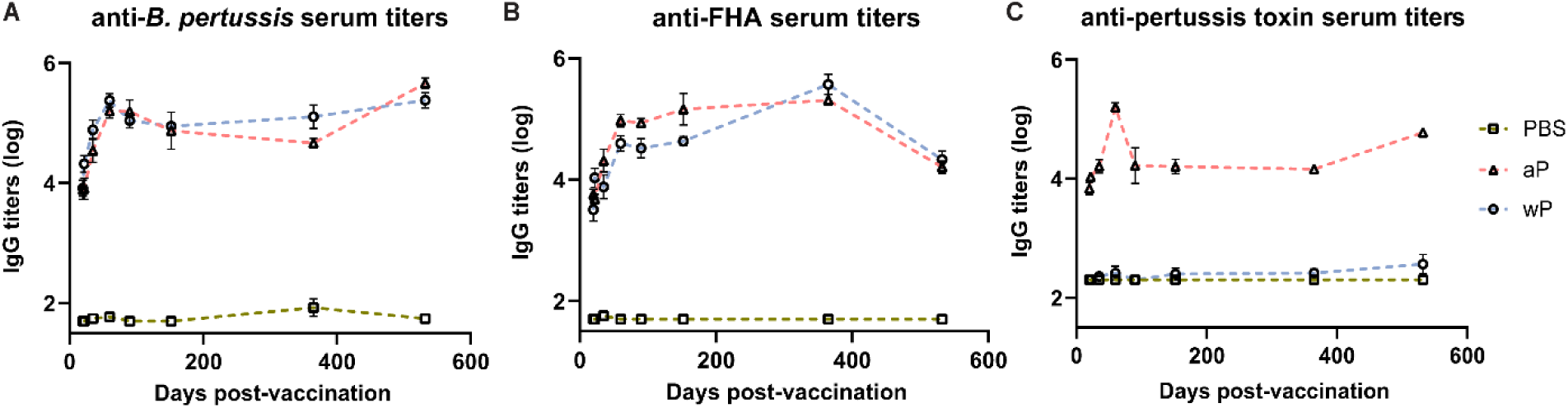
Pertussis specific antibody responses peak after boost and persist over time in immunized CD1 mice. Measured in serum of non-challenged mice collected at day 20, day 22, day 35, day 60, day 90, day 152, day 365, and day 532 post-vaccination. **(A)** IgG anti-*B. pertussis* antibodies in vaccinated mice (n=4-16) **(B)** IgG anti-FHA antibodies in vaccinated mice (n=4-16) **(C)** IgG anti-PT antibodies in vaccinated mice (n=4-16) Antibody responses were at or below the limit of detection in PBS vaccinated mice. Data were transformed using Y=Log(Y). Statistical analysis and p values are shown in Table S1.

In addition to opsonizing antibodies, it is imperative that pertussis immunization stimulates the production of PT-neutralizing antibodies that can prevent symptoms, disease manifestation, and in the case of infants, death^58^. Anti-PT antibodies have been proposed as a correlate of protection against pertussis; however, this is highly debated^57^. PT is an AB_5_ exotoxin that plays a key role in the pathogenesis of pertussis by triggering ADP-ribosylation which inhibits G protein-coupled signaling^59–64^. PT activity leads to a decrease in airway macrophages, induction of leukocytosis, and modulation of adaptive immune responses to *B. pertussis*. Unlike opsonizing antibodies, significant anti-PT IgG antibody responses were only observed in aP vaccinated animals compared to both vehicle control and wP immunized mice (**Fig 2C, Table S1**). Anti-PT antibodies peaked at day 60 post vaccination and remained elevated until day 532. To note, no anti-PT IgG antibodies were detected in wP immunized animals, consistent with the lack of PT in the NIBSC wP formulation due to manufacturing practices (**Fig 2C**).

DTaP is a combination vaccine also containing diphtheria toxoid and tetanus toxoid. Unlike the immunity to pertussis conferred by DTaP and Tdap immunization, immunity against diphtheria and tetanus does not wane overtime^65^. We observed that aP immunization elicits significant anti-diphtheria toxoid (**Fig S1A**) and anti-tetanus toxoid (**Fig S1B**) IgG antibody production compared to vehicle control mice at all time points. The anti-diphtheria toxoid and anti-tetanus toxoid IgG responses were similar to anti-PT responses as they increased significantly after prime and boost and remained high out to day 532 post-vaccination. Overall, the data indicate that antibodies against the whole bacterium, FHA, PT, diphtheria, and tetanus peaked post-boost and remained stable throughout the course of the study.

### wP but not aP immunization induces significant T_FH_ cell and CXCL13 responses

The presence of antibodies over a long period of time indicates that *B. pertussis* antigen-specific plasmablasts were produced in response to vaccination, were alive, and likely being repopulated^66^. However, antibody titers themselves do not directly predict vaccine recall capacity^67^. Conversely, cells associated with GC activity, such as T_FH_ cells and MBCs, are critical populations that dictate recall capacity. GC reactions take place in secondary lymphoid organs such as the spleen and lymph nodes^68^. Initiation of GC formation and the development of immunological memory relies on T_FH_ cells. T_FH_ cells are crucial for the survival and proliferation of B cells within the GC, and ultimately affinity maturation of B cells^69,70^. Our objective was to quantify T_FH_ cells in the secondary lymphoid organs of immunized mice to better understand how wP versus aP immunization influences GC activity.

First, we measured T_FH_ cells (CD4^+^CD3e^+^CXCR5^+^PD-1^+^)^71–73^ in the draining lymph node and spleen of immunized mice at time points between day 20 and day 532 post-vaccination (**Table S2**). In the right inguinal lymph node (**Fig 3A**) immunization with wP induced significant T_FH_ responses compared to aP and PBS immunized mice 20 days post-vaccination. Although no significant differences were observed between aP and wP immunized animals at other time points, T_FH_ cells in the draining lymph node at day 35 and in the spleen at days 20 and 22 were more numerous in the wP group (**Fig 3B**). Consistent with previous studies, a greater number of T_FH_ cells was only detected at the earliest time point studied (**Fig 3A**).

**Figure 3.**
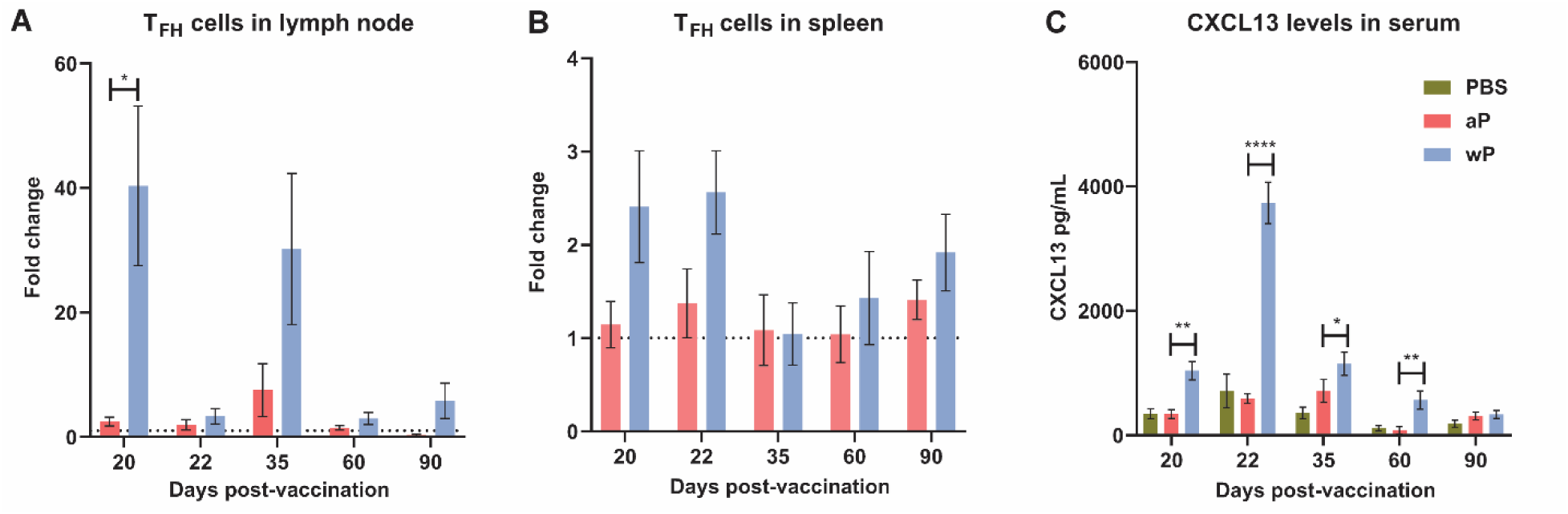
T follicular helper cell and CXCL13 responses are significant pre-boost in wP but not aP immunized CD1 mice. Flow cytometry was performed using single cell suspensions from both the **A)** draining lymph node and **B)** spleen to identify T_FH_ cells. **C)** CXCL13 levels (pg/mL) were measured in the blood serum of non-challenged, immunized mice. The *p*-values were calculated using an unpaired *t-*test, \**p* < 0.05, \*\**p* < 0.01, \*\*\**p* < 0.001, \*\*\*\**p* < 0.0001. Error bars are mean ± SEM values (n=4-16), aP (n=4-16), and wP (n=4-16).

To facilitate their function in the GCs, T_FH_ cells express the chemokine CXCL13 which is a signaling molecule that plays a crucial role in B cell recruitment and GC organization through binding to CXCR5^69^. While CXCL13 can be found locally in GCs it can also be measured in the serum as a biomarker of GC activity^45^. Therefore, we measured CXCL13 levels in the serum of immunized mice during the course of this experiment (**Fig 3C**). Levels of CXCL13 in the serum of vehicle control and aP-immunized mice were low (**Fig 3C**). In contrast, we observed that only wP immunization elicits significant production of CXCL13 compared to both aP and mock-immunized mice. CXCL13 levels in wP vaccinated animals peaked at day 22 post-vaccination and were significantly higher than aP mice at days 20 and 60 post-vaccination. Overall, the data suggest that T_FH_ cells and CXCL13 responses are differentially regulated in response to aP and wP early after vaccination, and that GC responses are greater in wP-vaccinated animals.

### wP immunization induces B. pertussis specific memory B cells in CD1 mice

MBCs are a vital component of the host’s immune system involved in protection against invading pathogens^74^. This population of cells is a product of the GC reaction and can be found in the spleen, lymph nodes, circulation, and more. MBCs are quiescent until recognition of antigen occurs. These cells can then rapidly respond by differentiating into plasma cells and mounting an antibody response. As MBCs are an important product of the GC reaction, and player in immunological memory, we sought to measure these cells over the course of this study in response to vaccination against *B. pertussis*.

To study MBCs, we incubated splenocytes with fluorescently labeled *B. pertussis*. We then separated MBCs from the rest of the splenocytes using a proprietary kit (Miltenyi), followed by labeling with CD3e, CD45R, IgG, CD38, and CD80^7,75^. We analyzed MBC populations based on labeling with *B. pertussis* (antigen-specific) and further defined the B cell populations based on CD38 and CD80 surface expression^75–77^(Fig **S2**). CD38 is an ectoenzyme with various functions and a transmembrane receptor in immune cells such as B cells^78^. CD38 is involved in B cell regulation and CD38 knockout mice exhibit deficiencies in antibody responses that result from T-cell-dependent antigens^78–80^. CD80 is a costimulatory molecule that plays a role in B and T cell interactions, and is expressed by both human and murine MBCs^81^.

We observed significant expansion of *B. pertussis*^+^ MBCs in wP immunized mice compared to both mock and aP immunized mice (**Fig 4A, Table S2, Fig S2**). In the wP group, this population peaked at day 35 post-boost, and although not statistically different from the mock vaccinated control group, *B. pertussis*^+^ MBCs were measurable at days 152 and 365 post-prime in wP vaccinated mice (**Fig 4A**, **Fig 4B**). We observed that in vaccinated animals, *B. pertussis*^+^ MBCs tend to be double positive for CD38^+^CD80^+^, while *B. pertussis*^-^ MBCs are mainly CD38^+^ (**Fig 4C, D, E, F**). This is likely important as IgG^+^ CD80^+^ MBCs have been shown to differentiate into antibody secreting cells with the capacity to seed GCs^82,83^. Interestingly, single-labeled CD80^+^ MBCs were low in both *B. pertussis*^+^ and *B. pertussis*^-^ populations. Overall, the data show that significant production of *B. pertussis*^+^ MBCs is most prevalent in wP immunized mice, and that *B. pertussis*^+^ MBCs are characterized by a unique combination of cell surface marker profiles.

**Figure 4.**
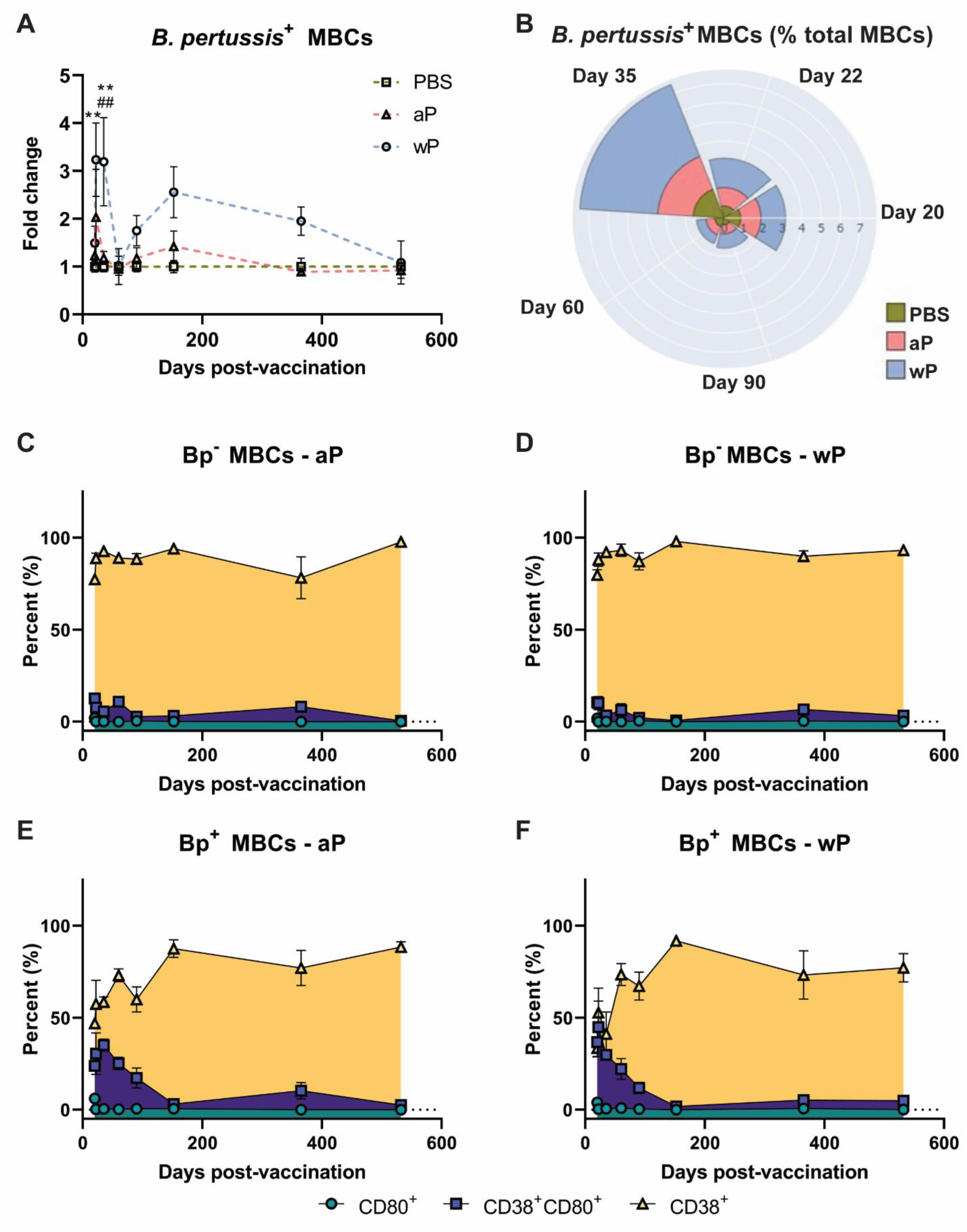
*B. pertussis*^+^ memory B cells are elevated in wP immunized CD1 mice and contribute to longevity of protection. Flow cytometry was performed using single cell suspensions from the spleen of immunized non-challenged mice. *B. pertussis*^+^ MBCs were identified following the protocol outlined in the methods. Extracellular markers were used to label *B. pertussis^+^* MBCs. **A**) Fold change of *B. pertussis^+^* MBCs in PBS, aP, and wP immunized animals. **B**) Percent total *B. pertussis^+^* MBCs. **Figures 4C, D, E,** and **F** show the percent of CD80^+^, CD38^+^CD80^+^, and CD80^+^ MBCs in immunized mice. **C**) *B. pertussis^-^* MBCs in aP immunized animals. **D**) *B. pertussis^-^* MBCs in wP immunized animals. **E**) *B. pertussis^+^* MBCs in aP immunized animals. **F**) *B. pertussis^+^* MBCs in wP immunized animals. The *p*-values were calculated using ANOVA followed by a Tukey’s multiple-comparison test, \*\**p* < 0.01. Error bars are mean ± SEM values (n=4-16), aP (n=4-16), and wP (n=4-16).

### Immunization against B. pertussis elicits production of antibody secreting cells in mice

In the GC, naïve B cells that receive T_FH_ cell help and undergo affinity maturation can differentiate into plasma cells. These cells migrate to the bone marrow and function to secrete antibodies and protect from infection. Long-lived plasma cells are terminally differentiated cells that can survive and continue to secrete antibodies for years, which has been demonstrated in both humans and mice^84^. We hypothesized that wP immunization would induce greater production of pertussis-specific long-lived plasma cells compared to aP immunization, mimicking the waning immunity observed in the human population. Therefore, we isolated bone marrow cells and quantified the number of total long-lived plasma cells and the number of antigen-specific antibody secreting cells.

Using flow cytometry, we first observed that there was no difference in the number of total long-lived plasma cells (CD19^-^CD45R^+^CD138^+^)^85^ in the bone marrow in any of the groups at any of the time points studied (data not shown). To determine the number of antigen-specific antibody secreting cells, we performed B cell ELISPOT on bone marrow samples (**Fig 5A-F**). We only observed a significant increase in the number of anti-*B. pertussis* antibody secreting cells in wP immunized mice one day post-boost compared to both mock and aP vaccinated animals (**Fig 5B**). Although there were no significant differences in antibody secreting cells at days 20 and 532 post-prime (**Fig 5A, C**), wP immunized animals had higher numbers of spots overall (**Fig 5A-C**). This contrasts with what was observed in the serological studies in which anti-*B. pertussis* titers remained high during the course of the study (**Fig 2A**).

**Figure 5.**
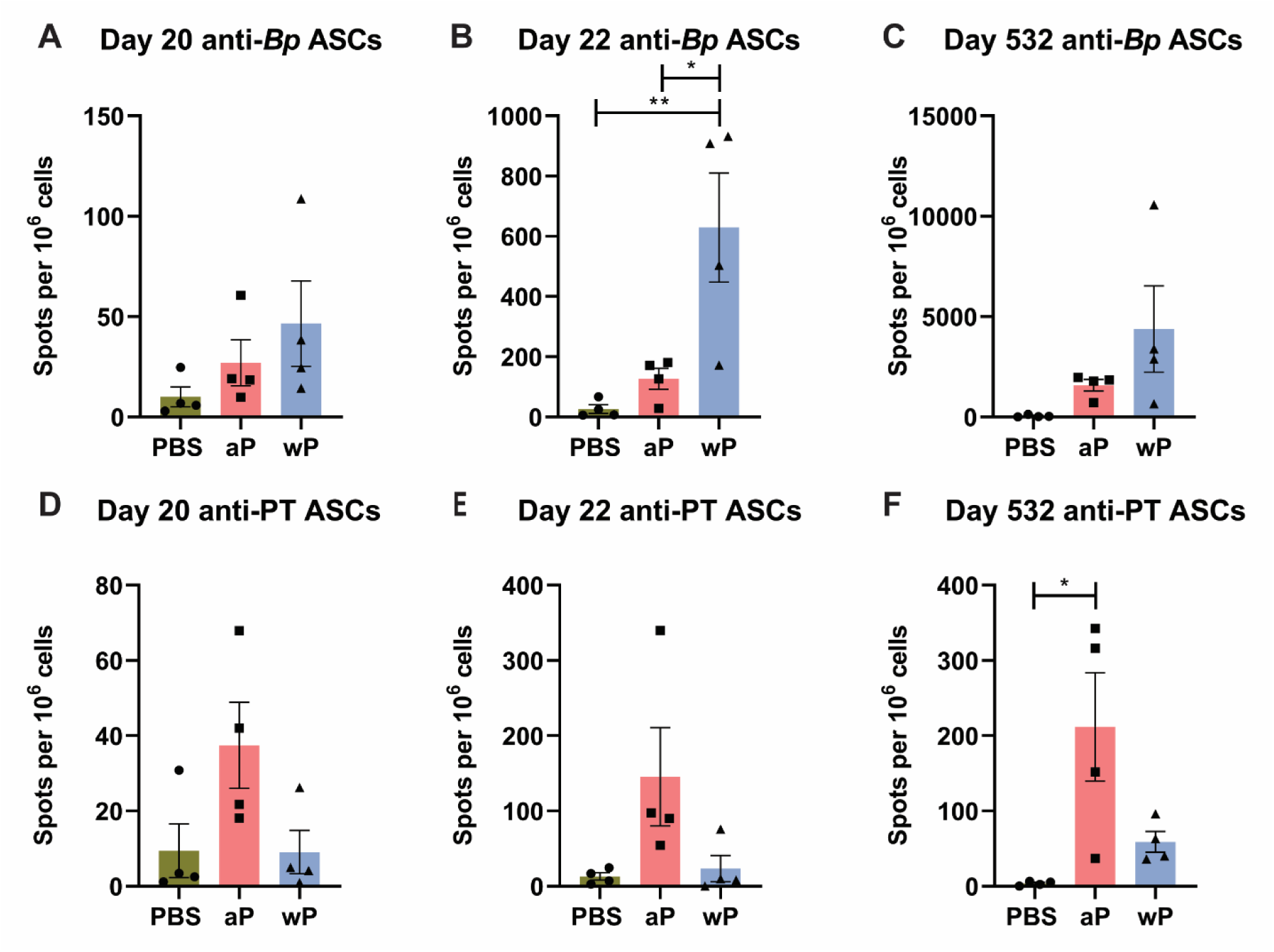
Anti-*B. pertussis* and anti-pertussis toxoid antibody secreting cells (ASCs) in immunized CD1 mice persist as far as day 532 post-prime. ASCs were analyzed at day 20, 22, and 532 post-vaccination in non-challenged, immunized mice. Anti-*B. pertussis* ASCs are shown for day 20 (**A**), day 22 (**B**), and day 532 (**C**). Anti-pertussis toxoid ASCs are shown for day 20 (**D**), day 22 (**E**), and day 532 (**F**). The *p*-values were calculated using mixed-effects analysis with a Tukey’s multiple-comparison test, \**p* < 0.05, \*\**p* < 0.01, for wP compared to PBS, *denotes comparison of wP to PBS and # denotes comparison of wP to aP groups. Error bars are mean ± SEM values (n=4), aP (n=4), and wP (n=4).

In this study, we also observed that anti-pertussis toxoid antibody secreting cells were present in greater numbers in aP mice compared to vehicle or wP mice, and persisted out to day 532 post-vaccination (**Fig 5F**). Notably, the number of anti-PT antibody secreting cells was significantly greater at day 532 post-prime in aP immunized animals (**Fig 5F**). This observation is consistent with the anti-PT titers detected in the serological study (**Fig 2C**) and with the fact that the wP vaccine used in this study contains minimal pertussis toxin. The data highlight important differences between serological detection of *B. pertussis* antibodies and quantification of antibody-producing cells in the bone marrow.

Altogether, the data described in this study highlight important differences in the immunological signatures triggered by aP and wP vaccination in mice (**Fig 6**). wP vaccination was characterized by stronger T_FH_ cell responses, CXCL13 production, *B. pertussis*^+^ MBCs, and anti-*B. pertussis* antibody secreting cells (**Fig 6**, **Fig 5**), compared to aP vaccination. These markers correlate with the stimulation of, or result from GC reactions, suggesting that wP vaccination triggers stronger follicular responses and vaccine-induced memory (**Fig 6B**).

**Figure 6.**
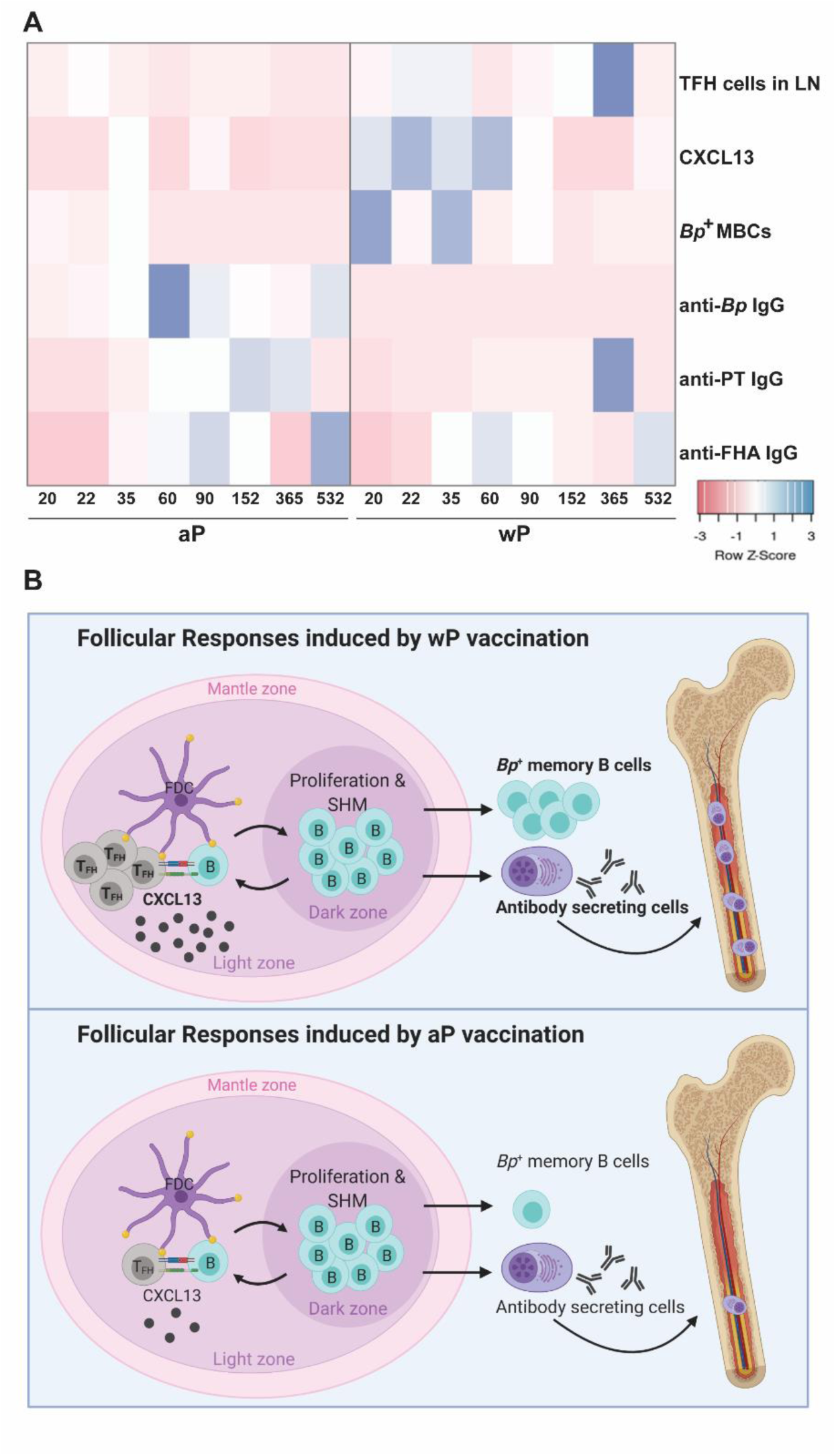
Predictors versus non-predictors of vaccine-induced memory. (**A**) Fold changes were calculated for each parameter and a heat map was generated (heatmapper.ca) comparing aP and wP non-challenged animals. (**B**) A model of vaccination and follicular responses highlighting differences in aP versus wP immunization.

## DISCUSSION

Pre-clinical animal models of vaccination and challenge have provided important insights into pertussis immunity. However, the majority of the recent conversations around pertussis vaccine immunity are focused on the pertussis T helper and T memory cell responses^57,86,87^. Furthermore, “longevity of protection” markers have not truly been identified in neither animals nor humans. Identifying biomarkers associated with vaccine-induced immunity that predict longevity of protection could bridge animal models and human vaccine trials, and help develop the next generation of pertussis vaccines. In the past, pertussis vaccine potency and efficacy were measured in pre-clinical models utilizing the Kendrick test, a lethal intracranial pertussis challenge model^88^. To replace the intracranial challenge model, lethal and sub-lethal aerosol and intranasal murine challenge models were utilized to measure pertussis vaccine efficacy^89–91^. The sub-lethal intranasal challenge model demonstrated statistical and reproducible differences in protection conferred by vaccination in short term experiments. These studies were helpful for determining that both cellular and humoral responses are involved in pertussis vaccine-mediated protection^89,92^. However, to this date, no absolute correlate of protection has been identified in the mouse model that can predict longevity of protection in humans.

During the development of acellular pertussis vaccines, immunogenicity of candidate vaccines was assessed in animals and humans^93^. Antibodies to PT, FHA, fimbriae, pertactin, DT, and TT were measured in immunized infants, along with toxin neutralization assays to determine levels of agglutinating antibodies^93^. Currently, immunoglobulin levels are measured and provide an approximation of vaccine efficacy; however, these metrics do not predict the duration of immunological memory and protection. Unfortunately, rarely in pertussis studies is the timing and longevity of protection considered due to the obvious challenges. This study addresses this gap in knowledge by determining long-term aP and wP vaccine-mediated protection out to day 532 post-vaccination, which we think is the longest-lasting pertussis vaccine study in mice performed to date. Laboratory shutdown during the COVID-19 pandemic increased our experimental time-frame but maybe for the better. In this study we used numerous approaches to characterize the follicular response to pertussis vaccination including antibody titers to vaccine antigens, CXCL13 levels in sera, T_FH_ cells, *B. pertussis* specific MBCs, and *B. pertussis*/PT specific bone marrow antibody secreting cells (**Fig 6**). We also identified serum levels of CXCL13 and *B. pertussis*-specific MBCs as potential biomarkers of pertussis vaccine-induced immune memory.

One of the challenges associated with monitoring the longevity of vaccine-mediated protection is that the model used must remain susceptible to infection over time. This was illustrated with the infant baboon model, which allows vaccine schedules to be studied and recapitulates human disease, but in which adults are not susceptible to pertussis infection (∼15 months of age). Therefore, we first studied the susceptibility of mice to *B. pertussis* over time. We assessed bacterial burden in the respiratory system of mice (lungs, trachea, and nasal wash) and found that susceptibility appears to change overtime, as seen in humans (**Fig 1B, C, D**). Neonates and unvaccinated infants are highly susceptible to pertussis infection; however, susceptibility decreases as they age toward adulthood. Furthermore, adults over 65 years of age are more likely to be hospitalized for pertussis than younger adults^94,95^. We observed a similar pattern of susceptibility in our vehicle control mice, in which mice between days 35 and 90 post-vaccination were susceptible to infection, but bacterial burden was below the limit of detection by day 152 post-vaccination. Bacterial burden was again detectable by day 365 post-vaccination.

One potential explanation behind the fluctuation in susceptibility to *B. pertussis* in mice, is that we suspect that like pigs, mice have differential expression over time of some inhibitory factors such as beta-defensin 1 that may render mice no longer susceptible to *B. pertussis*^96,97^. Alternatively, specific microbiota in the airway might out compete the challenge dose^98^. These data highlight the importance of conducting these types of studies over a long period of time, as intermediate lengths of studies may not allow measurement of vaccine-mediated efficacy due to low susceptibility. This is also important as susceptibility to *B. pertussis* in humans and in particular, death associated with pertussis, varies over time. Combinations of neonatal models with long-term models may be able to better evaluate pertussis vaccine efficacy. Additionally, inbred mice may have different windows of susceptibility that should be considered and studied further. Mahon *et al*. used BALB/c mice in long-term studies and control animals were susceptible to challenge at both 6 and 44 weeks after primary immunization^99^. Wolf *et al* also conducted short-term and long-term studies using BALB/c inbred mice and found that mock-vaccinated mice were susceptible to infection at both day 35 and 7 months, 3 days after challenge^28^. It is unclear why susceptibility changes over time in CD1 mice, and additional studies are needed to determine the cause, and if this phenomenon is strain or gender-specific.

In humans, studies clearly show that antibody titers against pertussis decay over time^100,101^. For example, human serum titers against PT decay as quickly as 6-12 months after vaccination whereas anti-tetanus serum titers last up to 19 years^102,103^. Similar to humans, BALB/c mice immunized with a low dose of wP or aP elicited high serum IgG antibody titers that increased rapidly after vaccination, but were undetectable by 6-9 months^92,99^. However, in our model and at 1/10^th^ the human dose of the vaccines, we did not observe antibody decay over time. In fact, antibody levels peaked after boost and remained elevated out to day 532 post-vaccination (**Fig 2, Fig S1**). These observations could be due, in part, to the relatively high dose of vaccine used here, and correlate with the sustained protection provided by both aP and wP vaccines over time.

To further investigate the humoral immune response to *B. pertussis* vaccination, we measured *B. pertussis*^+^ MBCs in the spleen of immunized mice (**Fig 4, Fig S2**) and the presence of antibody-secreting cells in the bone marrow. We observed that wP immunization elicited significant *B. pertussis*-specific MBC responses compared to both PBS and aP immunization (**Fig 4A, B**). Additionally, CD38^+^CD80^+^ cells were associated with *B. pertussis^+^* MBCs but not *B. pertussis^-^* MBCs (**Fig 4C, D, E, and F**). The unique memory profile associated with *B. pertussis^+^* MBCs support the hypothesis that B memory exists on a spectrum and could influence vaccine-induced memory. Further studies are needed to elucidate the importance of the notable cell marker profile associated with *B. pertussis^+^* MBCs.

Detection of antigen-specific MBCs from the spleen and antibody secreting cells in the bone marrow provides insight into the differences between wP and aP vaccine-induced immunity. Unfortunately, the protocols described here to detect antigen-specific MBCs are time consuming and technically challenging due to the rarity of these populations. Therefore, the implementation of antigen specific MBC analysis for clinical evaluation of pertussis vaccines in humans is unlikely. In addition, detection of antibody secreting cells requires invasive procedures to obtain samples for analysis, making its implementation unfavorable at the clinical level.

To identify additional markers of vaccine-induced memory, we measured CXCL13 in immunized mice as it has previously been measured in sera from humans and would be more feasible to monitor in clinical studies^104^. CXCL13 levels in the serum of immunized (non-challenged) mice peaked one day post-boost in wP-, but not in aP-immunized mice (**Fig 3C**), again highlighting another difference between both vaccines (**Fig 6A**). CXCL13 levels were significant both pre-boost and post-boost as far as day 60 post-vaccination (**Fig 3C**). Additional studies are needed to monitor the production of CXCL13 early on after vaccine priming. There appears to be a narrow window of CXCL13 production in mice, likely consistent with GC formation in response to vaccination. CXCL13 is a reliable plasma biomarker of GC activity in both humans and nonhuman primates^45^. Therefore, measuring CXCL13 levels post-vaccination using a minimally invasive blood draw could be utilized during clinical studies when testing candidate pertussis vaccines in humans.

There are obvious caveats to using CXCL13 as a biomarker for the longevity of pertussis vaccine-induced memory that need to be considered when designing clinical trials. The first is that CXCL13 is not antigen-specific. Another caveat is that CXCL13 expression is altered in response to infection, in cancer, systemic lupus erythematosus, rheumatoid arthritis, and other diseases involving germinal center, or ectopic lymphoid structure formation^105,106^. This should be taken in account when establishing exclusion criteria for clinical trials.

From this work, we propose that CXCL13, circulating *B. pertussis^+^* MBCs, and pertussis specific antibody titers should be measured together in the blood of patients enrolled in clinical trial vaccine trials to inform the longevity of vaccine-mediated protection (**Fig 6**). Future studies need to address if follicular responses induced by aP vaccination can be improved to levels similar to that induced by wP vaccination. Addition of adjuvants known to stimulate GC formation could enhance the longevity of aP vaccines and prevent waning immunity. This work strongly suggests that GC quantification, size, and composition should be evaluated in response to vaccination to identify formulations of the next generation pertussis vaccine that provide long-term protection.

## ACKNOWLEDGEMENTS

The study was supported by a 1R01AI14167101A1 to M.B.; K.L.W. received funding from the Cell and Molecular Biology and Biomedical Engineering Training Program funded by NIGMS grant T32 GM133369 (2019-2021), as well as the NASA West Virginia Space Grant Consortium Graduate Research Fellowship Program, Grant #80NSSC20M0055 (2021-2022). F.H.D is supported by NIH through grants 1R01AI137155-01A1 and 1R01AI153250-01A1. The WVU Vaccine Development Center, M.B. and F.H.D are supported by a Research Challenge Grant No. HEPC.dsr.18.6 from the Division of Science and Research, WV Higher Education Policy Commission. Flow Cytometry experiments were performed in the West Virginia University Flow Cytometry & Single Cell Core Facility, which is supported by the National Institutes of Health equipment grant number S10OD016165 and the Institutional Development Awards (IDeA) from the National Institute of General Medical Sciences of the National Institutes of Health under grant numbers P30GM121322 (TME CoBRE) and P20GM103434 (INBRE). The authors would like to thank Dr. Kathleen Brundage, director of the WVU Flow Cytometry and Single Cell Core Facility for her support.

---

**Figure S1.**
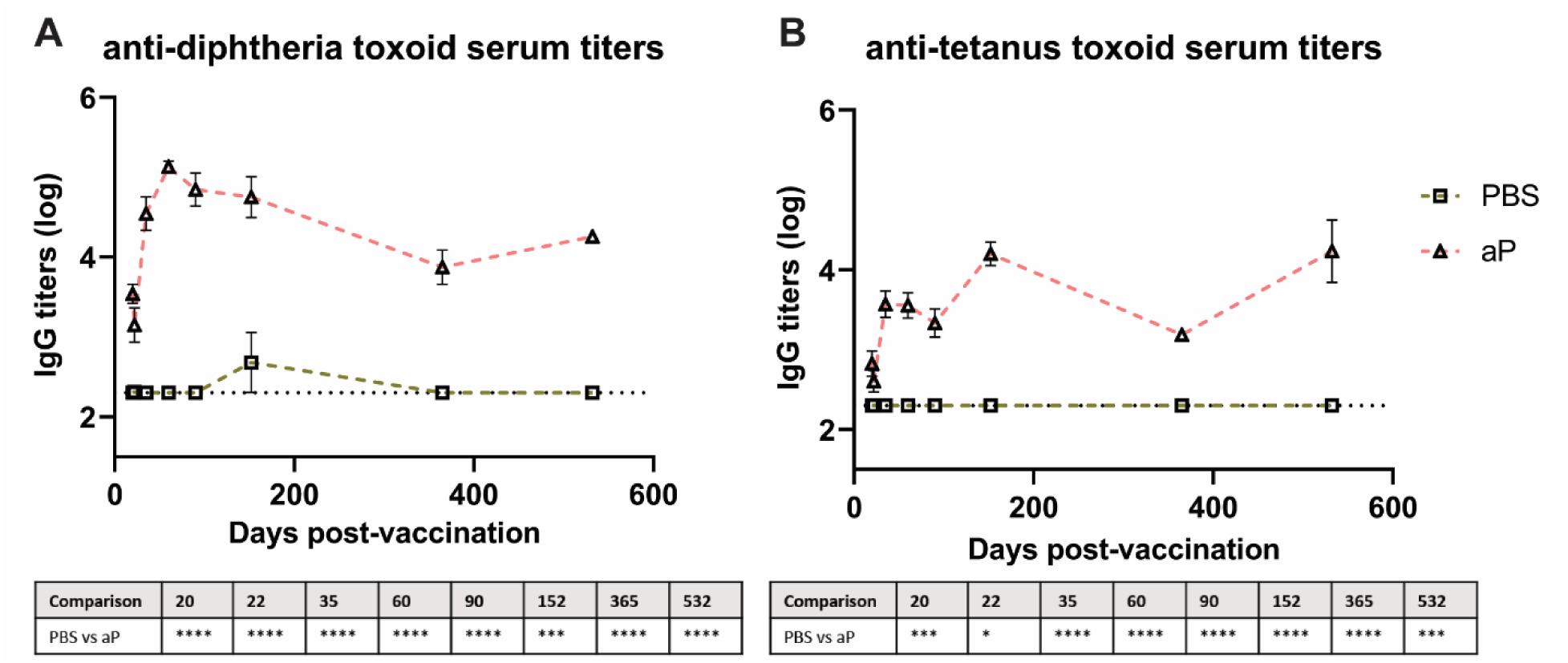
Anti-diphtheria and anti-tetanus antibodies persist as far as day 532 post-prime without waning. **(D)** IgG anti-diphtheria toxoid antibodies in non-challenged, vaccinated mice (n=4-16) and **(E)** IgG anti-tetanus toxoid antibodies measured in blood serum collected at day 20, day 22, day 35, day 60, day 90, day 152, day 365, and day 532 post-vaccination. Antibody responses were at or below the limit of detection in PBS vaccinated mice. Data were transformed using Y=Log(Y). The *p*-values were calculated using ANOVA followed by a Tukey’s multiple-comparison test, \**p* < 0.05, \*\*\**p* < 0.001, \*\*\*\**p* < 0.0001. Error bars are mean ± SEM values (n=4-16), aP (n=4-16), and wP (n=4-16).

**Figure S2.**
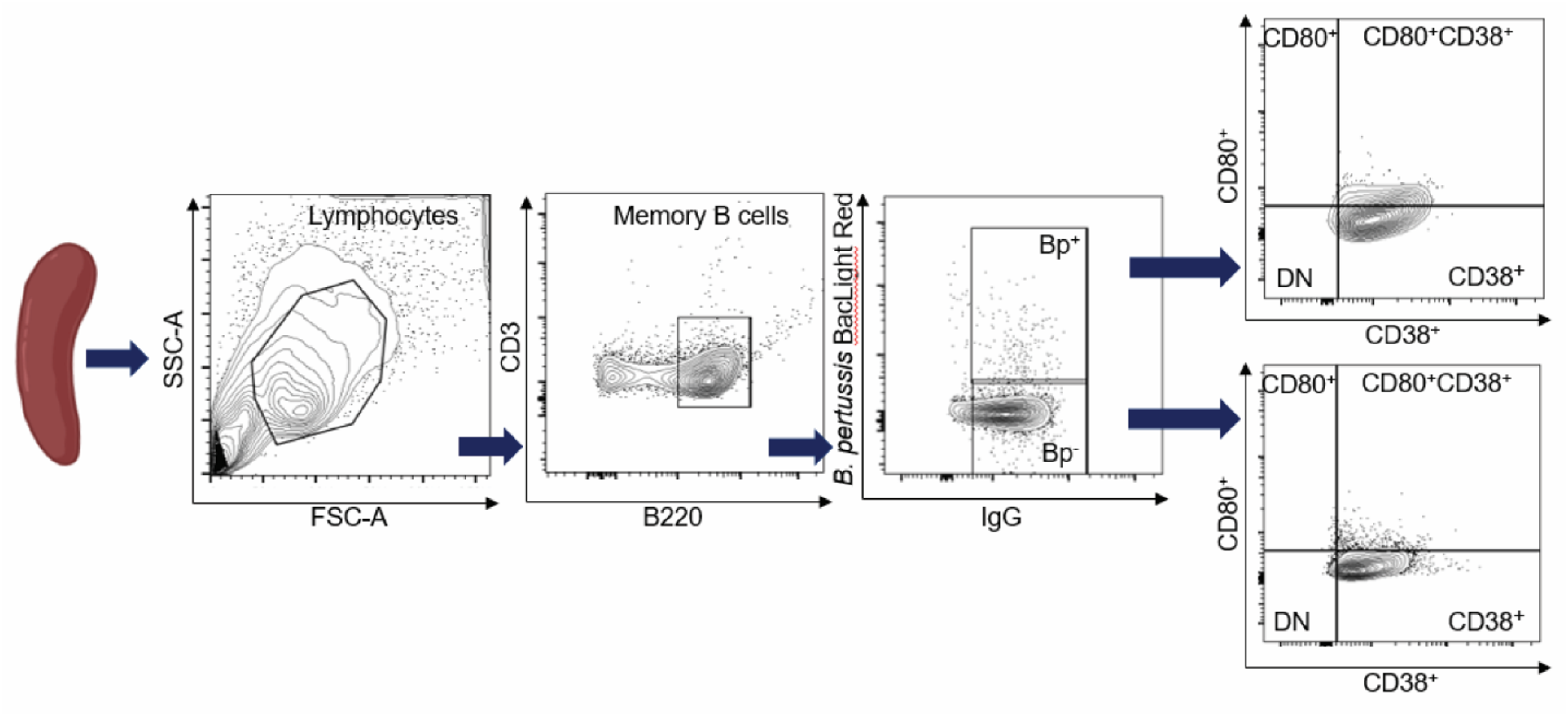
Flow cytometry allows for detection of *B. pertussis*^+^ MBCs. After selection of MBCs using our protocol and Miltenyi memory B cell isolation kit reagents, single live cells were gated to select for lymphocytes. From the lymphocyte population gating selected for CD3^-^ B220^+^ (CD45R) cells. From the B220^+^ (CD45R) population we gated to select *B. pertussis*^+^ cells ultimately isolating the *B. pertussis*^+^ MBC population. We further analyzed this population looking at CD38^+^, CD80^+^, CD38^+^CD80^+^ and double negative populations.

**Table S1.** Immunized mice have significantly elevated serum titer levels compared to mock-vaccinated animals. Statistics shown for **(A)** anti-*B. pertussis* serum titers **(B)** anti-FHA serum titers, **(C)** and anti-pertussis toxin serum titers. The *p*-values were calculated using ANOVA followed by a Tukey’s multiple-comparison test, *p < 0.05, **p < 0.01, ***p < 0.001, ****p < 0.0001. Error bars are mean ± SEM values (n=4-16), aP (n=4-16), and wP (n=4-16).

|  |  |  |  |  |  |  |  |  |  |
| --- | --- | --- | --- | --- | --- | --- | --- | --- | --- |
| **A** | **Comparison** | **20** | **22** | **35** | **60** | **90** | **152** | **365** | **532** |
|  | PBS vs aP | **** | **** | **** | **** | **** | **** | **** | **** |
|  | PBS vs wP | **** | **** | **** | **** | **** | **** | **** | **** |
|  | aP vs wP | NS | NS | NS | NS | NS | NS | NS | NS |
| **B** | **Comparison** | **20** | **22** | **35** | **60** | **90** | **152** | **365** | **532** |
|  | PBS vs aP | **** | **** | **** | **** | **** | **** | **** | **** |
|  | PBS vs wP | **** | **** | **** | **** | **** | **** | **** | **** |
|  | aP vs wP | NS | NS | NS | * | * | NS | NS | NS |
| **C** | **Comparison** | **20** | **22** | **35** | **60** | **90** | **152** | **365** | **532** |
|  | PBS vs aP | **** | **** | **** | **** | **** | **** | **** | **** |
|  | PBS vs wP | NS | NS | NS | NS | NS | NS | NS | NS |
|  | aP vs wP | **** | **** | **** | **** | **** | **** | **** | **** |

**Table S2.**
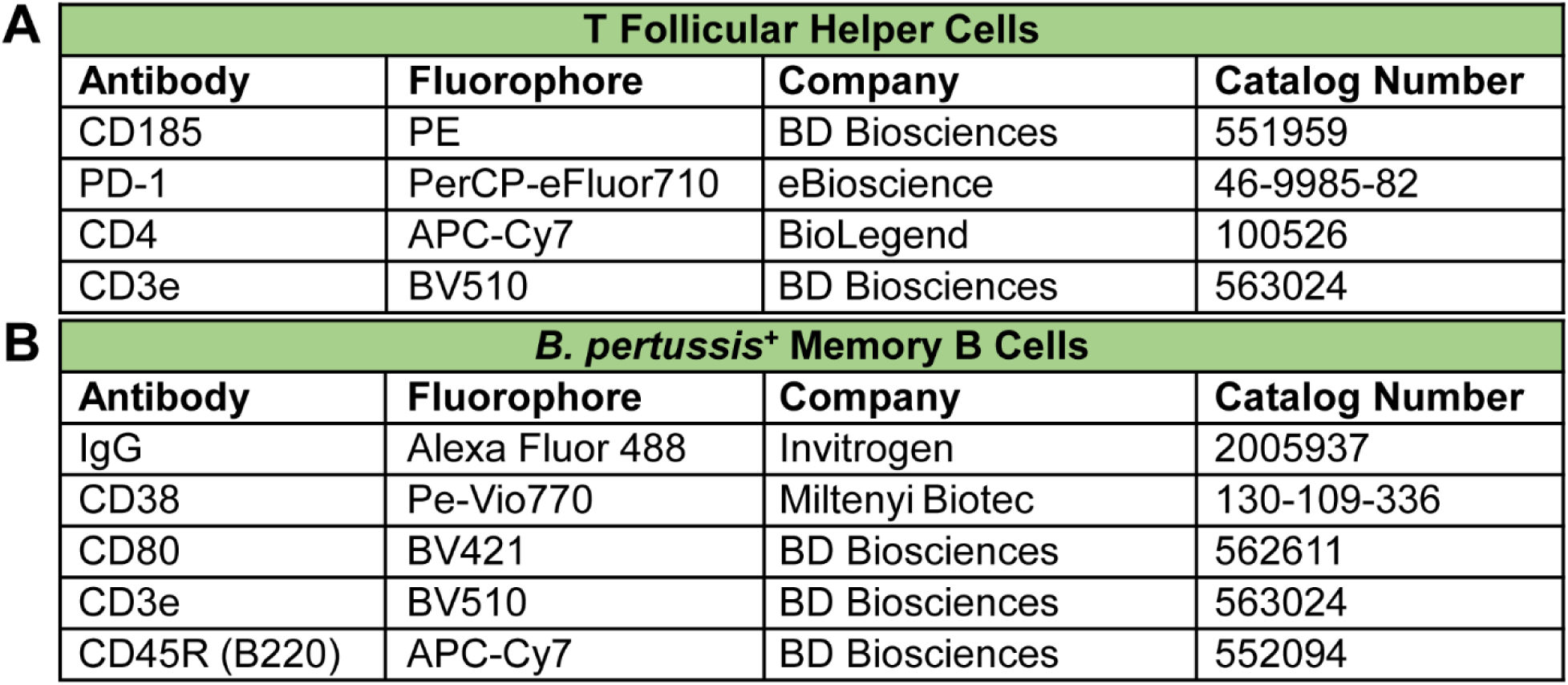
Flow cytometry panel design (**A**) T_FH_ cells and (**B**) *B. pertussis*^+^ MBCs.

